# Conserved Master Regulators Orchestrate Cellular Reprogramming-Induced Rejuvenation

**DOI:** 10.1101/2025.11.27.690899

**Authors:** A. Doğa Yücel, Harlan P. Stevens, Alexander Tyshkovskiy, Vadim N. Gladyshev

## Abstract

Partial somatic cell reprogramming has been proposed as a rejuvenation strategy, yet the regulatory architecture orchestrating age reversal remains unclear. Here, we performed gene regulatory network reconstruction across several independent systems to identify master regulators that coordinate reprogramming-induced rejuvenation (RIR). In mouse mesenchymal stem cells, mouse adipocytes, and human fibroblasts undergoing partial reprogramming, we identified genes showing opposite expression dynamics during aging and reprogramming. This approach revealed regulators governing rejuvenation rather than developmental programs. Despite divergent overall network architectures, nine transcription factors converged as master regulators across all three systems, including *Ezh2*, *Parp1*, and *Brca1*. These regulators undergo coordinated reorganization during reprogramming, characterized by broader target engagement and enhanced regulatory coherence. We further demonstrated that direct perturbation of *Ezh2* bidirectionally modulates transcriptomic age. Notably, overexpression of a catalytically inactive *Ezh2* mutant achieved rejuvenation, suggesting mechanisms distinct from canonical H3K27me3-mediated regulation are involved in RIR. Our findings reveal that cellular rejuvenation is orchestrated by conserved master regulators whose network coordination can be targeted independently of the reprogramming process.

## Introduction

Aging involves global changes across multiple hallmarks simultaneously.^1^ Efforts to tackle this process have mainly consisted of targeting specific hallmarks in the hope that fixing a hallmark would lead to improvements to the overall system. This approach has struggled to produce meaningful interventions for aging. ^2,3^ In contrast, cell reprogramming and, more recently, partial cell reprogramming have emerged as striking demonstrations that age-related changes could be targeted systemically.^4^ Unlike traditional aging interventions that act through stress responses or metabolic remodeling,^5^ these reprogramming approaches directly target the regulatory layer.^6–9^ This makes it a uniquely powerful system for dissecting the molecular basis of age reversal.

Studies on partial reprogramming have demonstrated that aged mammalian cells retain the capacity to reacquire youthful molecular and functional properties by transient relaxation of identity. It was reported that cyclic OSK(M) expression *in vivo* can restore vision, improve regeneration, reverse multiple transcriptional and epigenetic aging signatures, extend homogenous/heterogenous progeroid mice lifespan, and support wild-type mouse lifespan without teratoma formation.^4,10–15^ *In vitro*, partial reprogramming rejuvenates human fibroblasts and chondrocytes across transcriptomic, chromatin, and metabolic dimensions, producing widespread restoration of youthful regulatory features despite incomplete dedifferentiation and limited activation of pluripotency programs.^16,17^ These observations suggest that reprogramming-induced rejuvenation (RIR) ^18^ is separable from full cell fate reversal and can be achieved through targeted remodeling of the regulatory architecture that maintains core cell identity.

The contrast between full iPSC reprogramming and direct lineage conversion underscores this point. Fully reprogrammed iPSC-derived cells lose age-associated signatures, including DNA methylation patterns and transcriptional drift, consistent with global erasure of cis-regulatory memory.^19–21^ In contrast, induced neurons generated directly from fibroblasts retain donor-specific aging signatures, including transcriptional variability and epigenetic marks associated with drift.^21^ This divergence suggests that rejuvenation requires access to specific chromatin-maintenance mechanisms rather than lineage replacement alone. Partial reprogramming engages these mechanisms while preserving identity, whereas full reprogramming erases them and re-establishes new identities from a ground state.

These findings raise a fundamental question: what molecular systems allow partial relaxation of identity to restore youthful regulatory function while avoiding dedifferentiation? Previous work has identified chromatin regulators as central to this process. DNA methyltransferases Tet1 and Tet2 may be required for reprogramming-induced rejuvenation,^10^ and reprogramming-induced rejuvenated cells exhibit restored nucleosome regularity and recalibrated histone modification balance.^4^ However, identifying genes that change during rejuvenation does not reveal which factors actively drive the process versus those that respond as downstream consequences. Distinguishing upstream regulators from effector genes requires network-level analysis that can infer causal regulatory relationships.

Among these systems, Polycomb Repressive Complex 2 (PRC2) stands out mechanistically. PRC2 governs repression of developmental regulators,^22,23^ enforces identity boundaries,^20,22,24,25^ and stabilizes the epigenetic landscape through feedback-rich, bistable regulatory modules.^20,26^ Its catalytic subunits, EZH2 and EZH1, operate in distinct temporal regimes.^27^ EZH2 predominates in proliferative or early developmental contexts, rapidly depositing H3K27me3 to silence lineage-specifying modules.^28,29^ EZH1 predominates in quiescent or long-lived adult tissues, where it maintains chromatin compaction and low transcriptional responsiveness with slower kinetics.^30^ PRC2 subcomplex composition, recruitment logic, and antagonistic interactions with H3K4me3 and H3K36me2/3 establish threshold-dependent regulatory states that are stable yet flexible enough to support controlled identity transitions.^20,31^ These properties allow PRC2 to function not merely as a repressor but as a molecular system for cis-acting memory, integrating local chromatin marks, transcriptional activity, and architectural constraints.

With aging, PRC2-mediated regulation becomes imprecise. Developmental targets lose H3K27me3 fidelity,^32^ bivalent promoters become unstable,^33^ alternative lineage modules become aberrantly accessible,^34,35^ and broad heterochromatin regions accumulate diffuse H3K27me3 independent of canonical targeting.^36^ These patterns reflect a shift from coordinated repression to misallocated activity: a breakdown in the key epigenetic modulators that preserves identity. Epigenetic changes in PRC2 associated domains is one of the most consistent molecular signatures of aging across mammals,^37^ and age-dependent DNA methylation gain occurs predominantly in PRC2-bound low-methylated regions.^38^ This suggests that regulatory imbalance in PRC2 function is a conserved driver of age-related transcriptional instability.

In this study, we use cross-species, single-cell transcriptomic trajectories ^39,40^ and gene regulatory network reconstruction with NetID ^41^ to identify conserved transcriptional regulators that coordinate rejuvenation, decoupled from inducing pluripotency. Similar approaches were previously utilized to identify master regulators of key biological processes such as meiosis.^42^ By restricting analysis to genes whose expression change in opposite directions during aging and reprogramming, we isolate the regulators that specifically govern rejuvenation rather than developmental programs. Across mouse mesenchymal stem cells, mouse adipocytes, and human fibroblasts, we uncover strong convergence on chromatin- and architecture-regulating factors, with EZH2 emerging as a central node. Network-level changes reveal coordinated expansion of regulatory influence, increased coherence among master regulators, and systematic reorganization of chromatin-linked control.

Together, these findings show that partial reprogramming restores youthful cellular states by re-establishing the molecular machinery responsible for cis-acting memory and identity maintenance, rather than by reactivating pluripotency. They position EZH2 and associated chromatin regulators as core components of the rejuvenation process, linking the mechanistic basis of aging to the controllable properties of epigenetic maintenance systems.

## Results

### Network-level analysis identifies master regulators of reprogramming-induced rejuvenation

Building on the previously identified transcriptomic signatures of reprogramming and aging,^43,44^ we sought to uncover the regulatory architecture underlying reprogramming-induced rejuvenation (RIR). To identify master regulators of RIR, we performed gene regulatory network (GRN) reconstruction using NetID with Granger causality inference (L=30, maximum lagged time steps) ^41^ on single-cell transcriptomes (not used to construct the reprogramming signature) from three model systems: mouse mesenchymal stem cells (MSCs), mouse adipocytes, and human fibroblasts undergoing partial reprogramming^39,40^. For each system, we analyzed trajectories from baseline to reprogramming states in young (2–4 mo mouse; 22 y human) and aged (20–24 mo mouse; 96 y human) donors. We hypothesized that genes with significant opposite expression dynamics during reprogramming and aging represent direct targets of rejuvenation. Thus, we restricted GRN reconstruction to these “rejuvenation” genes (adjusted p < 0.05), ensuring our networks captured rejuvenation-associated regulatory interactions rather than general reprogramming or pluripotency programs (Fig. 1A). We refer to the modulation of such genes when we use the term rejuvenation in the GRN section of the paper.

**Figure 1.**
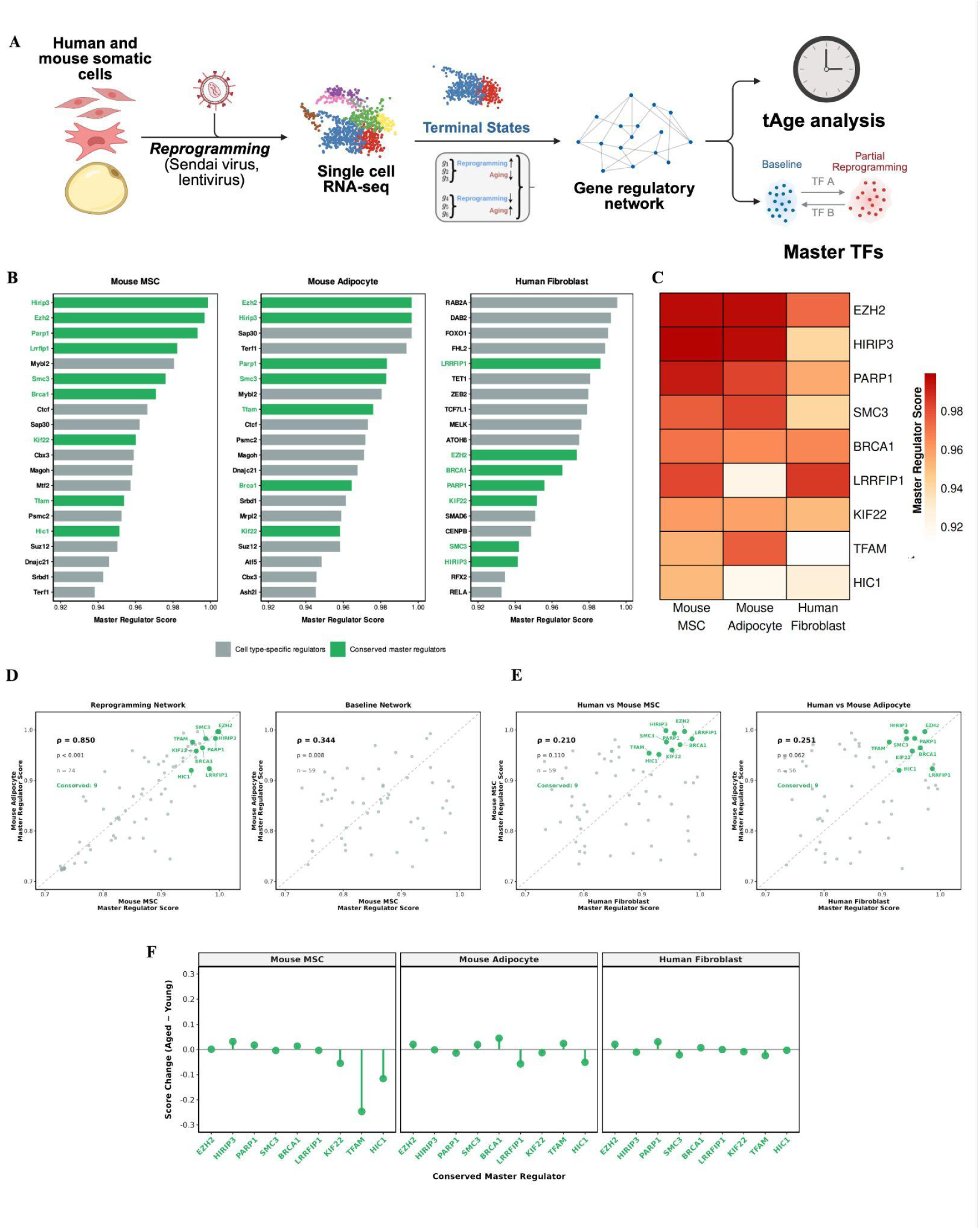
Gene regulatory network analysis identifies conserved master regulators of reprogramming-induced rejuvenation (RIR) **(A)** Schematic overview of the analytical pipeline. Single-cell RNA-seq data from human and mouse somatic cells undergoing partial reprogramming (Sendai virus, lentivirus) were analyzed to reconstruct gene regulatory networks using Granger causality inference. Master regulators were identified by integrating regulatory weight and eigenvector centrality. Transcriptomic age (tAge) was calculated for each cell along the reprogramming trajectory to assess cellular rejuvenation. **(B)** Top-ranked master regulators in mouse MSCs (left), mouse adipocytes (middle), and human fibroblasts (right). Bars show integrated master regulator scores (50:50 combination of regulatory weight and eigenvector centrality) for the top 20 transcription factors in each system. Green bars indicate conserved master regulators present in the top 30% across all three systems (n=9 factors). Gray bars indicate cell type-specific regulators. **(C)** Heatmap showing master regulator scores for the nine conserved factors across all three reprogramming systems. All nine factors exhibit high scores (>0.90) in all systems, with EZH2 showing particularly high scores (>0.97). Color intensity represents the master regulator score from 0.90 (white) to 1.00 (dark red). Missing values (white cells) indicate the factor was not detected in that system’s top 100. **(D)** Correlation of master regulator rankings within and between mouse mesenchymal lineages. Left: MSC vs. Adipocyte reprogramming networks show high correlation (Spearman ρ=0.850, 95% CI [0.744, 0.915], p < 0.001, n=74 shared TFs), with the nine conserved factors highlighted in green. Right: Baseline aging networks show substantially lower correlation (ρ=0.344, 95% CI [0.105, 0.547], p=0.008, n=59 shared TFs). The increase in regulatory coordination during reprogramming (Δρ=0.506) demonstrates activation of a shared rejuvenation program distinct from normal aging. The dashed line indicates identity (y=x). **(E)** Cross-species comparison of master regulator rankings. Human fibroblast reprogramming networks show lower overall correlation with mouse systems (Human vs. MSC: ρ=0.210, p=0.110, n=59 shared TFs; Human vs. Adipocyte: ρ=0.251, p=0.062, n=56 shared TFs), consistent with evolutionary divergence in regulatory network architecture. Despite low network-level correlation, the nine conserved factors (green) cluster in the upper right quadrant, indicating convergence on specific master regulators. The observed overlap of 9 factors in the top 30 across all systems is highly significant (permutation test: observed=9, expected=1.01 ± 0.96 SD, Z-score=8.31, p < 0.0001). **(F)** Master regulator score changes between young and aged cells during reprogramming. Each point represents the difference in master regulator score (Aged − Young) for one of the nine conserved factors. Master regulator hierarchies show high correlation between young and aged donors across all systems (MSC: ρ=0.890; Adipocyte: ρ=0.912; Human: ρ=0.842; all p < 0.001), indicating that the same transcription factors are recruited for rejuvenation regardless of starting cellular age. This consistency supports the universality of the rejuvenation regulatory program.

We identified master regulators by integrating network topology metrics: regulatory weight (direct transcriptional influence) and eigenvector centrality (network-wide influence) in equal proportions (Fig. 1B, C). Systematic comparison of six centrality metrics revealed that regulatory weight and eigenvector centrality provide complementary information (ρ ≈ 0.1-0.3). Closeness centrality, while also showing low redundancy, is poorly suited for sparse directed networks where many node pairs lack connecting paths. PageRank, Betweenness, and Degree showed high mutual correlations (ρ > 0.8), indicating redundancy (Supplementary Fig. 2D). This low redundancy between weight and eigenvector centrality justifies their integration for master regulator scoring.

Master regulator rankings between MSC and adipocyte reprogramming trajectories exhibited high correlation (Spearman ρ=0.850, 95% CI [0.744, 0.915], p < 0.001, n=74 shared TFs present in top-100 list of both systems), substantially higher than the correlation observed between their baseline aging trajectories (ρ=0.344, 95% CI [0.105, 0.547], p=0.008, n=59 shared TFs). This increase in regulatory coordination (Δρ=0.506, 95% CI excludes zero) indicates that mesenchymal lineages activate a shared regulatory program during rejuvenation that is distinct from their normal aging process (Fig. 1D). Having established convergence within mouse mesenchymal lineages, we next examined cross-species conservation. Cross-species comparisons revealed lower overall network correlation (human-mouse ρ=0.21-0.25), consistent with evolutionary divergence in regulatory network architecture (Fig. 1E). However, despite this network-level divergence, permutation analysis identified nine transcription factors that significantly converged across all three reprogramming systems. From a universe of 163 transcription factors (all TFs appearing in any system’s top-100 list), the following consistently ranked in the top 30 across all three systems: *Ezh2*, *Hirip3*, *Parp1*, *Brca1*, *Smc3*, *Lrrfip1*, *Kif22*, *Tfam*, and *Hic1* (permutation test: observed=9, expected=1.01 ± 0.96 SD, Z-score=8.31, p < 0.0001) (Fig. 1C, D, E) These factors showed high centrality scores across all systems (scores > 0.90), with Ezh2 ranking particularly high (scores > 0.97 in all three systems, from 0-1).

This pattern of low network-level correlation yet significant convergence on specific regulators suggests that while regulatory network architectures have diverged substantially between species, reprogramming-induced rejuvenation functionally converges on a core set of master regulators. The low overall correlation also indicates that our network method is not biased toward selecting the same transcription factors regardless of input, supporting the biological significance of the observed convergence. This is consistent with network rewiring that maintains functional constraints through conserved hub nodes.

Beyond their high absolute rankings, the nine conserved factors showed significantly larger master score increases during reprogramming compared to background transcription factors across all three systems (MSC: p = 0.022; Adipocyte: p = 0.025; Human: p < 0.001; Mann-Whitney U test; Supplementary Fig. 2B, C). This indicates that these factors are not just constitutive hubs but increase their positional importance in the network during the rejuvenation process, consistent with their functional role in orchestrating cellular state transitions.

Within each system, master regulators exhibited highly correlated rankings between young and aged cells during reprogramming (MSC: ρ=0.890, 95% CI [0.876, 0.902]; Adipocyte: ρ=0.912, 95% CI [0.902, 0.921]; Human Fibroblast: ρ=0.842, 95% CI [0.822, 0.860]; all p < 0.001), consistent with previous findings that mean iPSC reprogramming efficiency is comparable across age groups.^45^ The stability of master regulator hierarchies across donor ages suggests that rejuvenation engages the same core set of transcription factors regardless of the cell’s initial age (Fig. 1F).

### Core master regulators drive transcriptomic age reversal

Having established that reprogramming reorganizes regulatory networks into a more coherent architecture, we next asked whether master regulator activity influences cellular age. We measured transcriptomic age (tAge)^46^ across the reprogramming trajectory in the three model systems and found that reprogramming induced continuous transcriptomic rejuvenation across all systems. Both young and aged cells exhibited progressive decreases in tAge along the reprogramming trajectory (Spearman ρ=-0.53 to -0.90, all p < 0.001), with aged cells consistently displaying steeper anti-aging slopes than their young counterparts. This pattern held across mouse MSCs, mouse adipocytes, and human fibroblasts, indicating that partial reprogramming actively reverses age-associated transcriptional states rather than merely halting their progression (Fig. 2A).

**Figure 2.**
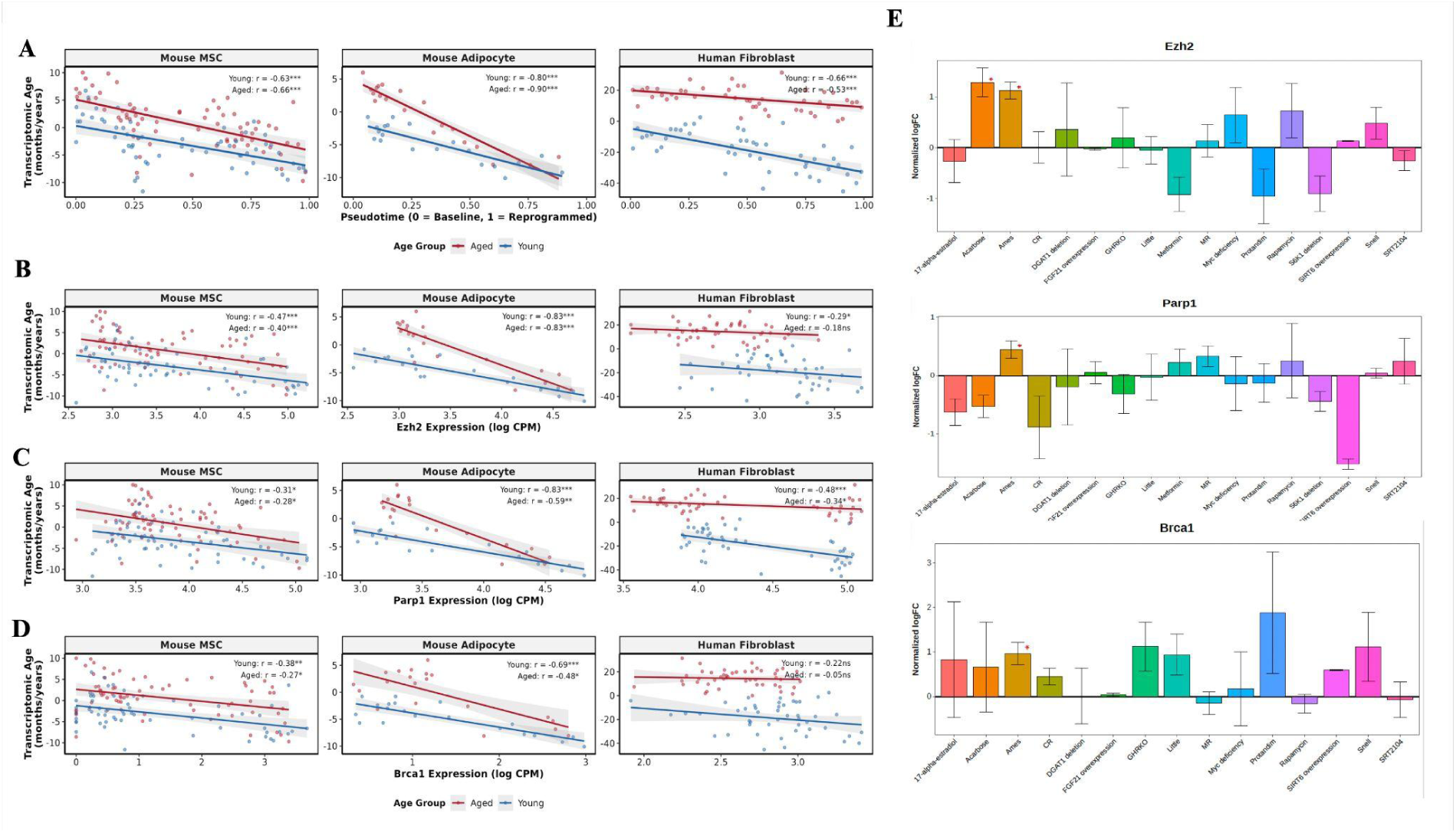
Core master regulators track transcriptomic age reversal. **(A)** Transcriptomic age decreases progressively along the reprogramming trajectory across all three systems. Scatter plots show relative transcriptomic age estimated with chronological multi-species multi-tissue clocks (months for mice, years for humans) versus pseudotime (0=baseline, 1=reprogrammed) for young (blue) and aged (red) cells. Both age groups show significant inverse correlations (all p < 0.001), with aged cells consistently displaying steeper rejuvenation slopes. Within each cell type, gene expression profiles were centered around the median of all cells. Linear regression lines with 95% confidence intervals (shaded regions) are shown. Correlation coefficients displayed in each panel. **(B)** EZH2 expression inversely correlates with transcriptomic age across all systems. Scatter plots show EZH2 expression (log CPM) versus transcriptomic age for young (blue) and aged (red) cells. Linear regression lines with 95% confidence intervals are shown. **(C)** PARP1 expression inversely correlates with transcriptomic age. Similar to EZH2, PARP1 shows strongest inverse correlations in mice. PARP1 demonstrates the most consistent inverse relationship with transcriptomic age across cell types. **(D)** BRCA1 expression shows cell-type-specific inverse correlations with transcriptomic age. Mouse adipocytes maintain strong inverse correlations, indicating cell-type-specific regulatory patterns. **(E)** Ezh2, Parp1 and Brca1 have higher baseline expression levels in Ames mice, the longest living strain of laboratory mice. * p < 0.05

To test whether master regulator expression directly correlates with cellular age reversal, we focused on *Ezh2, Parp1, and Brca1*, three of our top-ranked master regulators that have been causally implicated in the reprogramming process through previous gain and loss-of-function studies. ^47–50^ We examined the relationship between their expression levels and transcriptomic age across the reprogramming trajectory. All three master regulators showed inverse correlations with tAge, with expression levels rising as cells became transcriptionally younger. The strength and consistency of these associations varied substantially by cell type and regulator. Mouse adipocytes exhibited the strongest inverse correlations across all three factors (*Ezh2*: ρ=-0.83, *Parp1*: ρ=-0.83, *Brca1*: ρ=-0.69 in young cells; similar values in aged cells, all p < 0.001), with remarkably tight relationships between master regulator expression and transcriptomic age. Mouse MSCs showed moderate but consistent inverse correlations (ρ=-0.28 to -0.47), while human fibroblasts displayed more heterogeneous patterns, with *Parp1* showing the most robust associations consistent with Fig. 1 (young: ρ=-0.48, p < 0.001; aged: ρ=-0.34, p < 0.05) and *Brca1* exhibiting weaker, age-dependent relationships (young: ρ=-0.22, ns; aged: ρ=-0.05, ns).

Importantly, the inverse relationship between master regulator expression and tAge was present in both young and aged cells for most comparisons, suggesting that these factors are functionally involved in transcriptomic rejuvenation rather than merely marking cells that happen to be transcriptionally young. The cell-type-specific variation in correlation strength may reflect differences in baseline regulatory network architecture, epigenetic plasticity, or the specific reprogramming protocols used for each system. Nevertheless, the presence of significant inverse correlations across multiple master regulators, cell types, and age groups provides correlative evidence that master regulator activity is coupled to transcriptomic age reversal (Fig. 2B, C, D). Supporting the functional relevance of these master regulators, *Ezh2, Parp1, and Brca1* show significantly higher baseline expression in Ames dwarf mice compared to wild-type controls (Fig. 2E), suggesting that these factors may contribute to the longevity observed in this exceptionally long-lived mouse strain.^51^

### Reprogramming reorganizes networks for rejuvenation

Having identified master regulators, we next asked whether partial reprogramming alters not only the activity of individual regulators but also their collective coordination. We compared transcription factor regulatory weight correlations between baseline and partial reprogramming states in mouse MSCs, mouse adipocytes, and human fibroblasts. Across all young lineages, reprogramming led to a consistent increase in the effective out-degree (ΔDeff=12.3 ± 3.7) (MSC: ΔDeff=13.6 ± 3.5, p<0.01; Adipocyte: ΔDeff=8.2 ± 3.5, p<0.01; Human Fibroblast: ΔDeff=15.2 ± 6.2, p<0.01; Wilcoxon signed-rank test, n=9 TFs per system). This was accompanied by a corresponding decrease in regulatory inequality (ΔGini=−0.26 ± 0.04) (MSC: ΔGini=−0.28 ± 0.07, p<0.01; Adipocyte: ΔGini=−0.22 ± 0.08, p<0.01; Human Fibroblast: ΔGini=−0.29 ± 0.12, p<0.01).

Among the nine conserved regulators, five showed particularly large gains in regulatory breadth across all systems: *Ezh2, Smc3, Kif22, Brca1,* and *Hirip3* (mean ΔDeff=13.2 ± 1.3, range: 11.8 to 15.1), with Smc3 exhibiting the largest increase (ΔDeff=15.1, reaching +22.1 in human fibroblasts) alongside the most substantial reduction in inequality (mean ΔGini=−0.32, ranging from −0.41 to −0.13 across systems). These changes reflect a transition from specialized to more uniformly engaged regulation (Fig. 3A). Age-stratified analysis revealed that this reorganization capacity varies by cell type, with human fibroblasts maintaining robust reorganization in aged donors while mouse cells showed attenuated responses (Supplementary Fig. 3).

**Figure 3.**
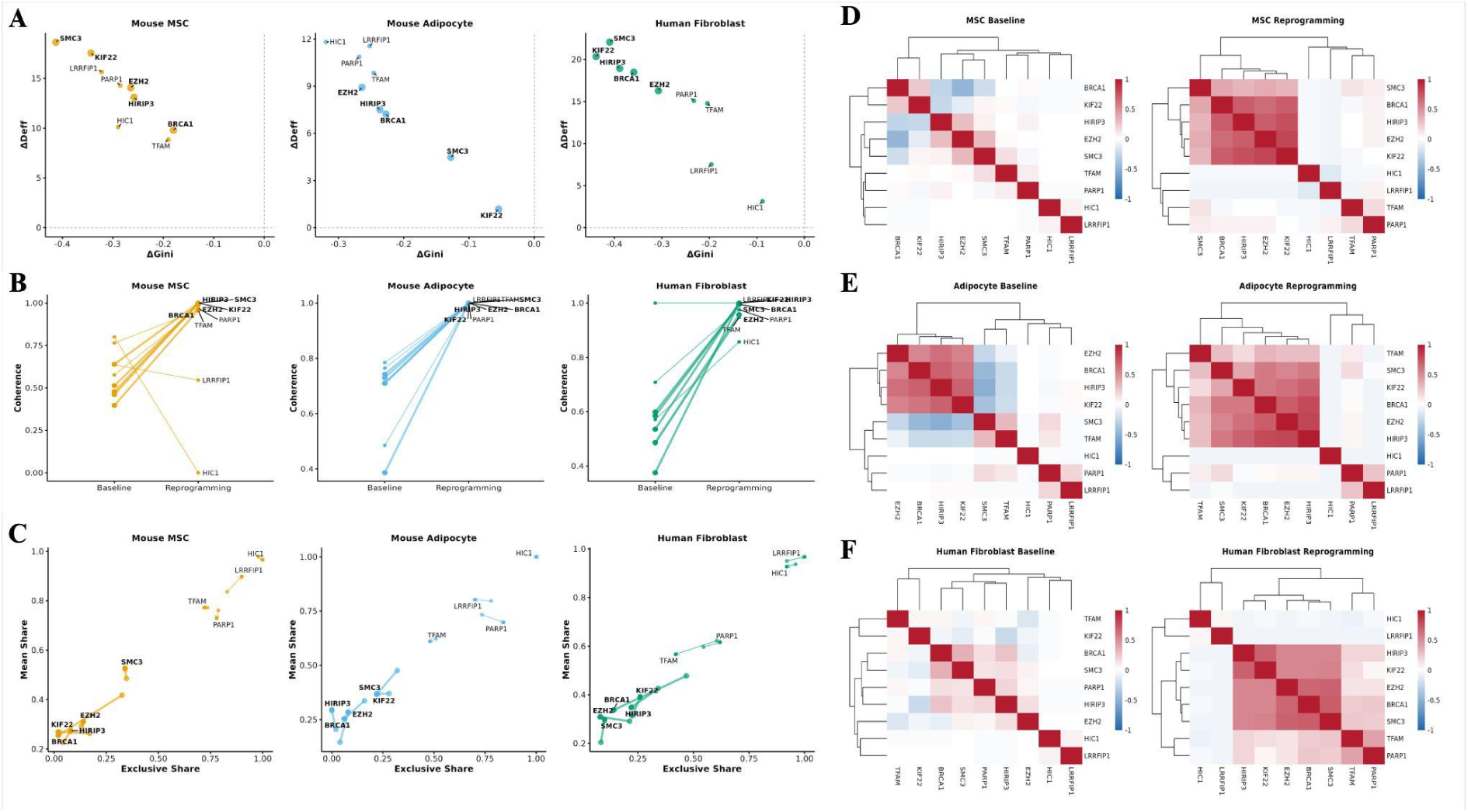
Network reorganization during reprogramming. **(A)** Changes in effective out-degree versus changes in Gini coefficient (ΔGini) for the nine conserved master regulators. Each point represents one transcription factor. Across all three systems, reprogramming increases effective out-degree (rightward shift, indicating broader target distribution) while decreasing Gini coefficient (downward shift, indicating more balanced regulation). The five core regulators (EZH2, SMC3, KIF22, BRCA1, HIRIP3) show particularly large shifts, labeled in each panel. Dashed lines indicate no change **(B)** Changes in regulatory coherence from baseline to reprogramming states. Lines connect the same transcription factor across states, with each line representing one of the nine conserved factors. Nearly all regulators show increased coherence during reprogramming (upward trajectories) across MSCs, adipocytes, and human fibroblasts, indicating enhanced coordination in regulatory direction among co-regulating transcription factors. The five core regulators are labeled at the reprogramming endpoint. **(C)** Regulatory weight distribution changes visualized as mean share versus exclusive share trajectories. Arrows show the transition from baseline (arrow origin) to reprogramming (arrowhead) for each conserved master regulator. Movement patterns indicate shifts from specialized, exclusive regulation toward more distributed, shared regulatory influence across all three systems. **(D)** Transcription factor (TF) regulatory weight correlation matrices for MSCs at baseline (left) and during reprogramming (right). Hierarchical clustering reveals the emergence of a tightly correlated subnetwork during reprogramming. Color intensity represents Spearman correlation coefficients from -1 (blue) to +1 (red). Overall mean pairwise correlation increases from ρ=0.00 at baseline to ρ=0.16 during reprogramming (Δρ=+0.16, p=0.11). The five core regulators (EZH2, BRCA1, HIRIP3, SMC3, KIF22) form a compact, highly correlated module in the reprogramming network, with mean correlation gains of +0.61 (from ρ=-0.03 to ρ=0.58). **(E)** TF weight correlation matrices for mouse adipocytes showing similar network reorganization. Overall mean pairwise correlation increases from ρ=0.06 at baseline to ρ=0.21 during reprogramming (Δρ=+0.15, p=0.09). The core five regulators show mean correlation gains of +0.42 (from ρ=0.21 to ρ=0.63). **(F)** TF weight correlation matrices for human fibroblasts. The core regulatory module emerges with the strongest reorganization among the three systems, with overall mean pairwise correlation increasing from ρ=0.03 to ρ=0.21 (Δρ=+0.18, p=0.002). Core regulator correlations increase by +0.55 (from ρ=0.05 to ρ=0.60).

Next, we quantified regulatory coherence, calculated as the degree to which transcription factors jointly activate or repress the same targets, and found substantial increases during reprogramming in most comparisons (overall ΔCoherence=0.32 ± 0.07). In MSCs and adipocytes, coherence rose for nearly all regulators (MSC: ΔCoherence=+0.24 ± 0.44, median=+0.39, p=0.16; Adipocyte: ΔCoherence=+0.34 ± 0.15, median=+0.28, p=0.014), with human fibroblasts showing the most pronounced increase (ΔCoherence=+0.38 ± 0.18, median=+0.41, p=0.014). The five core regulators exhibited particularly strong coherence gains (mean ΔCoherence=0.44 ± 0.03), indicating coordinated alignment toward shared transcriptional goals during rejuvenation (Fig. 3B).

At the network level, mean pairwise correlation among transcription factor regulatory weights increased in all three systems, though with varying statistical significance: ρ=0.00-0.16 in MSCs (Δρ=0.16, p=0.11), ρ=0.06 to 0.21 in adipocytes (Δρ=0.15, p=0.09), and ρ=0.03 to 0.21 in human fibroblasts (Δρ=0.18, p=0.002; paired Wilcoxon test, n=36 TF pairs per system). The increase was most prominent among the five core regulators (*Ezh2*, *Brca1*, *Hirip3*, *Smc3*, and *Kif22*) which transitioned from weak or negative mean correlations at baseline (MSC: ρ=−0.03; Adipocyte: ρ=0.21; Human Fibroblast: ρ=0.05) to strong positive correlations after reprogramming (MSC: ρ=0.58; Adipocyte: ρ=0.63; Human Fibroblast: ρ=0.60), representing mean correlation gains of +0.61 in MSCs, +0.42 in adipocytes, and +0.55 in fibroblasts. These master transcription factors, initially acting independently at baseline, became strongly co-regulated after reprogramming, indicating the formation of a compact chromatin and DNA repair-associated subnetwork, given the roles of these factors.^52–59^ (Fig. 3C-F).

Together, these results show that partial reprogramming reorganizes existing regulatory connections into a more synchronized and cooperative structure rather than establishing entirely new ones. This network reweighting creates a coordinated transcriptional program centered on chromatin organization, genome maintenance, and cell cycle control that operates across cell types and species.

### *Ezh2* Bidirectionally Controls Transcriptomic Age

To establish causal relationships between master regulator activity and cellular age, we analyzed data from inhibition and overexpression experiments focusing on *Ezh2*, the top-ranked master regulator. siRNA inhibition of *Ezh2* in human dermal fibroblasts ^60^ significantly increased transcriptomic age across both mortality (p < 0.01) and chronological (p < 0.05) clocks, demonstrating that Ezh2 activity is necessary for maintaining youthful transcriptomic states (Fig. 4A).

**Figure 4.**
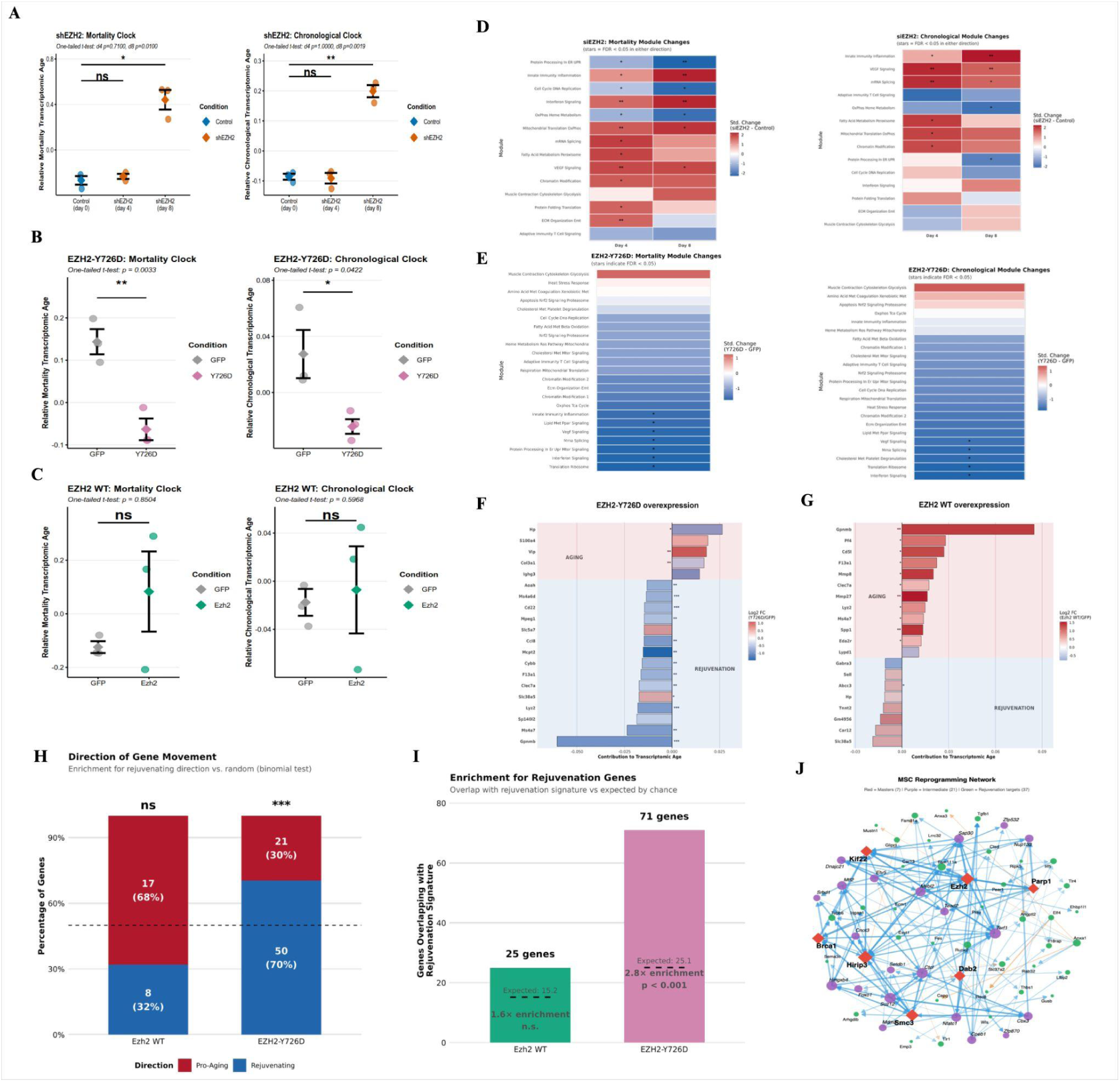
Ezh2 controls transcriptomic age through non-canonical mechanisms. **(A)** EZH2 inhibition accelerates transcriptomic aging in human fibroblasts (GSE109064). Left: Mortality multi-species multi-tissue aging clock shows significant increase in transcriptomic age following shEZH2 treatment at day 8 compared to day 0 controls (One-tailed t-test on day 0 ctrl vs day 8: p < 0.0033). Right: Chronological multi-species multi-tissue aging clock similarly shows progressive increase in transcriptomic age with EZH2 knockdown (One-tailed t-test on day 0 ctrl vs day 8: p < 0.0006). Points represent individual biological replicates (n=3 per condition); error bars show mean ± SEM. Significance: ** p < 0.01, *** p < 0.001. **(B)** Catalytically inactive EZH2-Y726D mutant overexpression in retinal ganglion cells reduces transcriptomic age (GSE247320). Left: Mouse multi-tissue mortality clock shows significant decrease in transcriptomic age with Y726D expression compared to GFP control (One-tailed t-test: p < 0.0033). Right: Mouse multi-tissue chronological clock shows similar rejuvenation effect (One-tailed t-test: p < 0.0422). Points represent individual biological replicates (n=3 per condition); error bars show mean ± SEM. **(C)** Wild-type EZH2 overexpression shows no significant effect on transcriptomic age in retinal ganglion cells (GSE247320). Neither mouse multi-tissue mortality clock (left) nor mouse multi-tissue chronological clock (right) show significant changes with EZH2 WT overexpression compared to GFP control (ns=not significant). This dissociation between WT and Y726D effects suggests EZH2 drives rejuvenation through mechanisms independent of its canonical PRC2-mediated H3K27me3 deposition. Points represent biological replicates; error bars show mean ± SEM. **(D)** EZH2 inhibition dysregulates mortality and chronological module-specific clocks. Heatmap shows changes in module-specific transcriptomic age (Δ tAge, standardized) at day 4 and day 8 following shEZH2 treatment relative to day 0 baseline. Red indicates age acceleration; white indicates no change. Modules span diverse aging hallmarks including innate immunity, interferon signaling, mitochondrial function, and ECM organization. Most modules show progressive age acceleration (red), consistent with overall aging phenotype. Asterisks indicate FDR-adjusted significance: * p < 0.05, ** p < 0.01. **(E)** EZH2-Y726D rejuvenates cells according to mortality and chronological module-specific clocks. Heatmap shows module-specific transcriptomic age changes with Y726D overexpression. Blue indicates age reversal (rejuvenation). Multiple aging-associated modules show significant rejuvenation, including innate immunity, lipid metabolism, and protein processing pathways, demonstrating that EZH2-Y726D achieves broad rejuvenation across multiple aging hallmarks simultaneously. Asterisks indicate significance: * p < 0.05. **(F)** Differential contributions of genes to chronological tAge change induced by EZH2-Y726D overexpression. Y726D overexpression shows coordinated changes across aging-associated and rejuvenating gene modules, with strong effects on multiple pathways. **(G)** Gene contribution to tAge change induced by wild-type EZH2 overexpression shows less coordinated effects with opposing contributions (both red and blue), potentially explaining the lack of net effect on overall transcriptomic age. Bar color intensity represents contribution magnitude and direction. **(H)** EZH2-Y726D selectively activates rejuvenation gene programs. Stacked bar chart shows directional concordance between differentially expressed genes (DEGs) and rejuvenation signatures. Wild-type EZH2 shows 32% of DEGs (8/25 genes) moving in the rejuvenating direction (blue), which is not significantly enriched above random expectation (binomial test: ns). In contrast, EZH2-Y726D shows 70% of DEGs (50/71 genes) moving in the rejuvenating direction, significantly enriched above random expectation (binomial test: p < 0.001, ***). The remainder move in the pro-aging direction (red). Numbers indicate gene counts and percentages. Asterisks indicate enrichment vs 50% random: *** p < 0.001, ns=not significant. **(I)** Overlap between differentially expressed genes and rejuvenation signature. The bar chart shows the number of genes from EZH2 overexpression experiments that overlap with the rejuvenation gene signature, compared to expected overlap by chance (dashed line). Wild-type EZH2 overexpression shows 25 genes overlapping (1.6× enrichment, ns), while EZH2-Y726D shows 71 genes overlapping (2.8× enrichment, p < 0.001, hypergeometric test), demonstrating that the catalytically inactive mutant specifically engages rejuvenation-associated gene programs. **(J)** Master regulators form a core hub in the MSC reprogramming network. Network diagram shows regulatory interactions among transcription factors, with master regulators (purple nodes) occupying central positions. Node size represents regulatory influence; edge color indicates regulatory direction (green=activation, cyan/blue=repression). The five core master regulators (EZH2, BRCA1, HIRIP3, SMC3, KIF22) are labeled and form a densely interconnected subnetwork. Green nodes indicate rejuvenation-associated transcription factors. This network architecture demonstrates that master regulators coordinate rejuvenation through a hierarchical regulatory cascade.

To test whether *Ezh2*’s rejuvenating effects require its canonical histone methyltransferase activity, we compared wild-type *Ezh2* overexpression to a catalytically inactive mutant (Y726D) in retinal ganglion cells.^61^ Surprisingly, wild-type overexpression showed no significant effect on tAge in either mortality-associated or chronological dimensions. In contrast, overexpression of the catalytically inactive Ezh2-Y726D mutant significantly reduced transcriptomic age in both aging dimensions (Fig. 4B, C).

This dissociation between wild-type and Y726D effects ^61^ suggests that *Ezh2* drives rejuvenation through mechanisms independent of its canonical PRC2-mediated H3K27me3 deposition. Beyond its role as the catalytic subunit of Polycomb Repressive Complex 2, *Ezh2* possesses alternative transcriptional regulatory functions, including direct transcription factor methylation and PRC2-independent chromatin remodeling.^62,63^ We hypothesize that wild-type overexpression may trigger compensatory stress responses or reach saturation of endogenous pathways, while Y726D specifically engages the non-canonical rejuvenation mechanism without these confounding effects. Consistent with this interpretation, module-level analysis revealed that *Ezh2*-Y726D affected multiple aging hallmarks like inflammation and mTOR signaling simultaneously, with significant effects observed across chronological aging modules and mortality-associated pathways (Fig. 4D, E). These findings establish a direct causal relationship between *Ezh2* activity and tAge, demonstrating that rejuvenation can be achieved by overexpressing specific master regulators.

### Master Regulator Targets Drive Rejuvenation Gene Expression Programs

At the individual gene level, top contributors to transcriptomic age changes such as *Gpnmb*, *Ms4a7*, and *Clec7a,* showed divergent responses to WT versus Y726D overexpression. WT *Ezh2* shifted these genes toward pro-aging expression patterns, whereas Y726D shifted them in the opposite, anti-aging direction (Fig. 4F, G). To determine whether master regulators achieve transcriptomic age reversal through coordinated regulation of downstream targets, we analyzed the overlap between genes differentially expressed following *Ezh2*-Y726D overexpression and previously defined rejuvenation gene signatures.^43^ Directional analysis revealed that 70% of shared genes moved toward the anti-aging direction following Ezh2-Y726D expression, significantly exceeding the 32% concordance observed with wild-type Ezh2 (Fisher’s exact test, p < 0.001) (Fig. 4H). Genes affected by *Ezh2*-Y726D showed significant enrichment for rejuvenation signatures (p < 0.001, hypergeometric test), with 71 genes overlapping between *Ezh2*-Y726D targets and expected rejuvenation genes (Fig. 4I). This indicates that Ezh2 mutant achieves transcriptomic age reversal by suppressing age-associated gene expression programs rather than triggering compensatory stress responses. Network analysis of master regulator targets revealed a hierarchical regulatory cascade, with master transcription factors regulating intermediate genes that subsequently control rejuvenation-associated target genes (Fig. 4J).

### Ezh2-Mutant Rejuvenation Signals Through Chromatin Accessibility Changes

To validate and better understand the rejuvenation signal seen in the transcriptome following *Ezh2*-Y726D overexpression, we analyzed the chromatin accessibility profiles of both the mutant Ezh2-Y726D and wild-type *Ezh2* overexpression. We analyzed differentially accessible regions (DAR) for each condition relative to their respective controls. When comparing each set of DARs to the rodent aging meta-signature, we observed a positive correlation for Ezh2-WT (r=0.21, p < 0.0001) and a negative correlation for *Ezh2*-Y726D (r=–0.21, p < 0.0001). This suggests that genes upregulated with age are more likely to become accessible in *Ezh2* overexpression and to become less accessible in *Ezh2*-Y726D overexpression and vice versa, providing a mechanistic link to the gene expression differences noted above (Fig. 5A). This pattern was strongest at promoter-associated DARs, consistent with promoter peaks more directly influencing gene expression (Fig. 5B).

**Figure 5.**
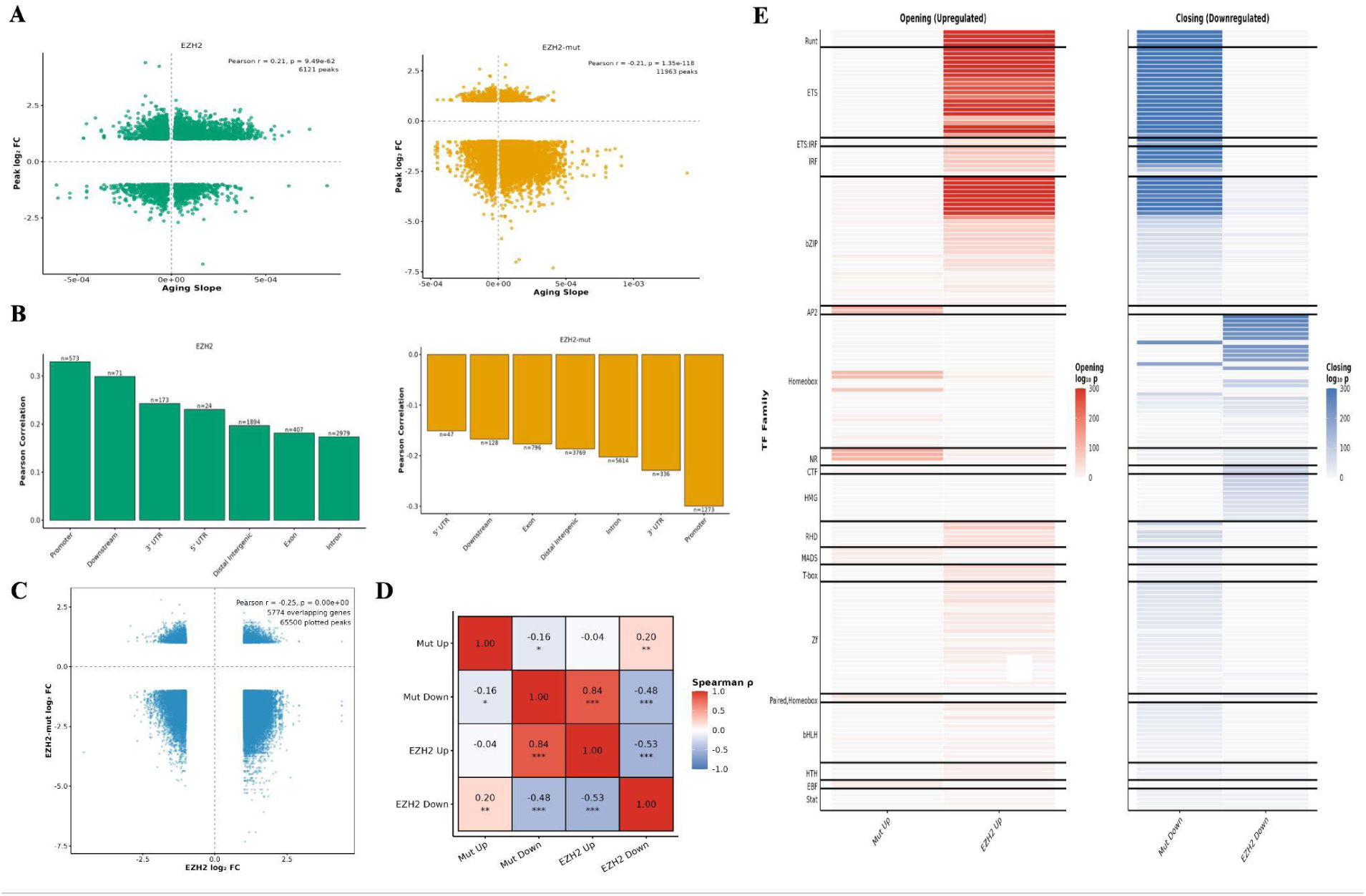
Ezh2 induces transcriptomic rejuvenation through chromatin accessibility. **(A)** Chromatin accessibility changes correlate with aging signatures in opposing directions. Volcano plots show the relationship between aging slope (x-axis) and chromatin accessibility log fold change (y-axis) for differentially accessible regions (DARs). Left: Wild-type EZH2 overexpression shows positive correlation (Pearson r=0.21, p < 0.0001), indicating that regions gaining accessibility are enriched for age-upregulated genes (pro-aging pattern). Right: EZH2-Y726D shows negative correlation (Pearson r=-0.21, p=1.13e-115), indicating that regions gaining accessibility are enriched for age-downregulated genes (anti-aging pattern). Green and orange points represent individual DARs; dashed lines indicate significance thresholds. **(B)** Genomic region-specific correlations between chromatin accessibility and aging signatures. Bar plots show Pearson correlations between DARs and aging slopes across different genomic features. Left: Wild-type EZH2 shows positive correlations across most regions, strongest in promoters (Pearson r > 0.30). Right: EZH2-Y726D shows negative correlations, with strongest anti-aging effects in promoter regions (Pearson r > -0.30). This demonstrates that the mutant reorganizes chromatin accessibility at functionally relevant regulatory regions to oppose aging-associated chromatin changes. Error bars represent correlation confidence intervals where applicable. **(C)** Chromatin accessibility changes show opposing patterns between EZH2-Y726D and wild-type at overlapping genes. Scatter plot shows the relationship between differentially accessible regions (DARs) for EZH2-Y726D (x-axis) versus EZH2-WT (y-axis) at genes with overlapping measurements (n = 5774 genes from 65500 ATAC-seq peaks). The moderate negative correlation (Pearson r = -0.25, p < 0.001) indicates that the catalytically inactive mutant and wild-type EZH2 have opposing effects on chromatin accessibility, consistent with their divergent effects on transcriptomic age. **(D)** Chromatin accessibility changes show opposing patterns between EZH2-Y726D and wild-type. The correlation matrix shows Spearman correlations between differentially accessible regions (DARs) in different conditions. EZH2-Y726D opening peaks show strong negative correlation with wild-type opening peaks (Spearman = -0.16, **), while showing positive correlation with wild-type closing peaks (Spearman = 0.20, **). This inverse relationship demonstrates that the catalytically inactive mutant reorganizes chromatin accessibility in the opposite direction from wild-type EZH2, consistent with their opposing effects on transcriptomic age. Asterisks indicate significance: ** p < 0.01, *** p < 0.001. **(E)** Transcription factor motif enrichment reveals distinct chromatin regulatory programs. Heatmaps show transcription factor binding motif enrichment in opening (upregulated, left) and closing (downregulated, right) chromatin regions for EZH2-Y726D mutant and wild-type overexpression. Color intensity represents log2 enrichment β coefficient. Major TF families show divergent patterns: Runt/Ets motifs are enriched in wild type opening but Y726D closing peaks, while Homeobox motifs show the opposite pattern. This demonstrates that EZH2-Y726D and wild-type recruit distinct transcriptional programs through differential chromatin remodeling, with Y726D favoring regulatory architectures associated with cellular rejuvenation.

Consistent with the original study,^61^ we found a moderate negative correlation of the DARs at overlapping genes between the mutant Ezh2-Y726D and wild-type Ezh2 (r=-0.25, p < 0.001), indicating opposing effects on chromatin accessibility (Fig. 5C, D).^61^ TF motif enrichment analysis of opening and closing peaks further underscored the divergence between the two conditions. Within each condition, motifs enriched in opening peaks were negatively correlated with those in closing peaks. Surprisingly, peaks that closed in *Ezh2*-Y726D overexpression were strongly correlated with those that opened in *Ezh2-*WT, whereas the inverse relationship was much weaker. At the transcription factor family level, this trend persisted: Runt and ETS motifs were enriched in *Ezh2*-WT opening and *Ezh2*-Y726D closing peaks, while Homeobox motifs were enriched in *Ezh2*-WT closing peaks and NR/AP2 motifs in *Ezh2*-Y726D opening peaks (Fig. 5E). The accessibility differences between the *Ezh2*-Y726D and *Ezh2*-WT support the transcriptomic findings and suggest mechanistic differences between the *Ezh2* forms that mediate their differential impact on rejuvenation.

## Discussion

We identified nine transcription factors that converge as master regulators of reprogramming-induced rejuvenation across three independent mammalian systems. During reprogramming, these regulators undergo coordinated network reorganization characterized by broader target engagement and enhanced regulatory coherence. This reorganization reflects the active process of cellular state transition, with master regulators adopting broader regulatory patterns that orchestrate changes across multiple subsystems. Overexpression of *Ezh2* mutant demonstrates that tAge reversal may operate through non-canonical mechanisms distinct from its pluripotency-associated functions, providing direct evidence that transcriptomic age reversal and developmental reprogramming can be mechanistically separated. Previous studies have shown that there is a strong convergence between rejuvenation and aging associated changes on PRC2 targets,^64^ and *Ezh2* overexpression was shown to reverse aging-associated changes.^65–67^ Our network-based approach not only supports these findings but also uncovers additional master regulators as candidates for multifactorial rejuvenation therapies.

The cross-species convergence on the same master regulators despite divergent overall network architectures suggests these nodes are under evolutionary constraint. This pattern is consistent with network evolution models where peripheral connections diverge while core functional hubs are maintained.^68^ The biological functions of these regulators provide insight into why they may be conserved. Rather than serving singular roles, many regulators exhibit multi-functional properties that bridge different cellular systems. *Ezh2* functions both as a chromatin regulator through PRC2 and in DNA repair through PARP1 methylation and DDB2 stabilization.^62,69^ Smc3, a part of the cohesin complex, organizes chromatin architecture while antagonizing or mediating polycomb interactions.^58,70–72^ *Parp1* links DNA repair with metabolic regulation and transcription.^73,74^ This multi-functionality may explain their importance as integration points, where single regulators coordinate changes across chromatin organization, genome maintenance, and metabolic networks.

The network reorganization we observed during reprogramming, with increased regulatory breadth and coherence, appears paradoxical at first. One might expect activated master regulators to show more focused, specialized regulation of their targets during reprogramming. Instead, they exhibit broader engagement across multiple targets. We propose that this reflects the fundamental requirements of coordinated cellular state transitions. Reprogramming requires simultaneous changes across chromatin structure, DNA repair capacity, metabolic state, and transcriptional programs.^50,75–78^ Notably, the capacity for this reorganization showed age-dependent variation in mouse cells, where aged donors exhibited attenuated network remodeling compared to young donors. Human fibroblasts, however, maintained reorganization capacity even in aged donors, suggesting species-specific differences in network plasticity that warrant further investigation. Achieving this coordination may require master regulators to temporarily adopt broader regulatory patterns that orchestrate changes across multiple subsystems. This interpretation is supported by the consistency of the pattern across different cell types undergoing the same transition process.

The *Ezh2* perturbation experiments provide mechanistic insight into how rejuvenation can be decoupled from pluripotency. The catalytically inactive Y726D mutant achieves superior rejuvenation compared to wild-type despite lacking the H3K27me3 methyltransferase activity ^79^ that is central to PRC2-mediated developmental regulation.^23,80^ This suggests *Ezh2* possesses alternative functions that drive transcriptomic age changes independently of its canonical role in maintaining pluripotency and silencing developmental genes. The divergent chromatin accessibility patterns between wild-type and Y726D further support distinct mechanisms of action. These findings align with our previous observation that only 35% of reprogramming-induced rejuvenation changes involve pluripotency-associated genes.^44^ The non-canonical functions of *Ezh2* may account for a substantial portion of the remaining 65%, offering a molecular explanation for how rejuvenation and pluripotency can be separated.

The multi-functional nature of master regulators has important implications for understanding aging mechanisms. Rather than viewing aging as failure of individual pathways, our findings suggest it may reflect progressive loss of coordination between interconnected cellular systems. Chromatin regulation, DNA repair, metabolic homeostasis, and transcriptional control normally operate in coordination, maintained by regulators that bridge these systems. During aging, this coordination may deteriorate, leading to subsystem-specific failures that compound over time. Rejuvenation, in this framework, represents restoration of coordination rather than simply reversing individual molecular changes. This perspective is consistent with the idea that epigenetic aging reflects loss of regulatory fidelity and that interventions work by restoring coherent regulation rather than targeting isolated defects.

The observation that *Tfam*, a mitochondrial transcription factor, appears among the convergent regulators is particularly interesting given recent findings that its deficiency in T cells causes systemic premature aging through inflammaging.^81^ This suggests some master regulators may exert effects beyond their cell of origin, potentially through secreted factors or cell-cell communication. Whether other identified regulators have similar systemic properties remains to be determined, but it raises the possibility that cellular rejuvenation could have organism-level consequences through both autonomous and non-autonomous mechanisms.

Our study has several limitations that warrant consideration. First, our network analysis employs Granger causality inference to identify directional regulatory relationships from single-cell trajectory data. While this computational approach provides evidence for causality through temporal precedence along pseudotime, it infers regulatory directionality rather than definitively establishing it through experimental perturbation. We partially address this limitation through testing the causal role of *Ezh2* perturbation on transcriptomic age regulation. While the causal role of *Ezh2* was demonstrated in this study, the mechanistic roles of the other eight regulators and their synergistic effects in rejuvenation remain to be directly tested. Second, we focused on transcriptional networks, but post-transcriptional regulation, protein-level changes, and metabolic alterations also contribute to cellular aging and may involve additional regulatory layers. Third, our analysis examined cells during active reprogramming, but whether network properties return to more focused patterns after transition completion is unknown. Time-course data extending beyond the reprogramming window would help distinguish transient coordination patterns from stable regulatory states. Fourth, while we tested the Ezh2 effect on molecular age in one system, the contributions of other master regulators and their potential redundancy across different cell types require systematic investigation.

Our findings suggest several directions for future research. The mechanism by which catalytically inactive *Ezh2*-Y726D achieves tAge reversal warrants detailed investigation. Does it work through PARP1 methylation, DDB2 stabilization, or other non-canonical functions? Are these mechanisms sufficient, or do they require coordination with other master regulators? Testing whether similar separations between canonical and non-canonical functions exist for other regulators could reveal general principles of transcriptomic regulation. The consistency of network reorganization patterns across cell types suggests that other cellular transitions could have similar dynamics and supports further research into whether differentiation, senescence entry, or tissue regeneration show comparable network dynamics. Understanding whether this represents a universal feature of cell state transitions versus a reprogramming-specific phenomenon would clarify its biological significance. Finally, the therapeutic potential of targeting non-canonical functions deserves exploration, though substantial work remains to translate these findings into practical interventions.

This work reveals the regulatory architecture underlying reprogramming-induced rejuvenation and demonstrates that age reversal can operate through mechanisms distinct from developmental programming. By identifying the master regulators that converge across human and mouse systems despite divergent network architectures and characterizing how their coordination patterns change during cellular state transitions, we provide a framework for understanding how transcriptomic age can be modulated independently of cell fate. These findings bridge our previous transcriptomic analysis of rejuvenation signatures with the regulatory logic that orchestrates those changes, moving toward a more complete mechanistic understanding of how reprogramming reverses aging-associated molecular damage.

## Experimental Procedures

### Data Acquisition and Preprocessing

Single-cell RNA-sequencing data for mouse mesenchymal stem cells (MSCs), mouse adipocytes, and human fibroblasts undergoing partial reprogramming were obtained from previously published studies.^39,40^ We used the processed and quality-controlled count matrices from the original publications to maintain consistency with their analyses and cell state annotations. For mouse datasets, we analyzed cells from young (2-4-month-old) and aged (20-24-month-old) donors. For human fibroblasts, we analyzed cells from young (22 years) and aged (96 years) donors. Cell state annotations (baseline, reprogramming, etc.) were retained from the original studies. Log-normalized expression values (logcounts) were used for all downstream analyses. D0 and D7 Fibroblast and Early Pluripotent

### Transcription Factor Identification

TFs were identified using the comprehensive mouse TF list from SCENIC (allTFs_mm.txt, downloaded from https://resources.aertslab.org/cistarget/tf_lists/) for mouse datasets, and the human TF list (allTFs_hg38.txt) for human fibroblasts. TFs were filtered to include only those present in the expressed gene set for each dataset.

### Aging and Reprogramming Signatures

Aging signatures were derived from our rodent aging meta-analysis.^43^ To identify genes specifically associated with rejuvenation rather than general reprogramming or pluripotency programs, we selected target genes showing opposite expression dynamics between reprogramming and aging. For each significant gene in both signatures, we determined its directional trend (upregulated: slope > 0; downregulated: slope < 0). Genes were classified as rejuvenation-associated if they showed opposite directional trends between the reprogramming and aging signatures (e.g., upregulated during reprogramming but downregulated during aging, or *vice versa*). For both signatures, genes were considered significant if they showed consistent directional changes (FDR-adjusted p < 0.05) and were present in the expressed gene sets. These were used as the target gene set for subsequent GRN reconstruction.

### Trajectory Inference

For each dataset (young and aged separately), we performed trajectory inference using the FateDynamic module of the NetID package (version 0.1.1). To ensure consistent trajectory orientation, we selected optimal start cells representing the most differentiated baseline state in each dataset. We calculated a reprogramming score for each cell as the mean expression of the top 50 rejuvenation-associated genes. The cell with the lowest reprogramming score within the baseline/adipogenic cluster was selected as the start cell.

Trajectory inference was performed using Palantir with the following parameters: start cell as defined above, terminal states set to baseline and reprogramming states for each cell type, number of highly variable genes=1000, minimum counts=10, and velocity integration disabled. This yielded pseudotime values ranging from 0 (baseline state) to 1 (reprogrammed state) for each cell, with pseudotime subsequently inverted (1 - pseudotime) to represent progression from baseline to reprogramming.

### Gene Regulatory Network Reconstruction

Gene regulatory networks were reconstructed using the NetID framework (version 0.1.1) with Granger causality inference. The complete NetID pipeline consisted of four steps:

Step 1: Initial NetID Run. We first ran NetID without dynamic inference to prepare the data structure and perform initial quality control using the RunNetID function. This step used geosketch for efficient sampling of the expression manifold with normalization disabled to preserve count-based variance structure.

Step 2: Trajectory Inference. As described above, FateDynamic was applied to infer developmental trajectories and calculate pseudotime values for each cell.

Step 3: NetID with Dynamic Inference. NetID was re-run with dynamic inference enabled using RunNetID with dynamicInfer=TRUE, incorporating the pseudotime information from Step 2. This step models gene expression dynamics along the trajectory and prepares data for causal inference.

Step 4: Causal Network Inference. Gene regulatory networks were inferred using FateCausal with Granger causality testing (L=30, representing a lag parameter of 30 for temporal dependency modeling). This step generates directed regulatory edges from transcription factors to target genes, with edge weights representing the strength and direction (positive/negative) of regulatory influence. Networks were constructed separately for each terminal state (e.g., adipogenic baseline vs. reprogramming state for adipocytes).

This complete pipeline was applied independently to young and aged cells for each of the three cell types (MSCs, adipocytes, human fibroblasts).

### Master Regulator Identification

To identify master regulators within each GRN, we integrated two complementary network topology metrics: (1) regulatory weight, quantified as the sum of absolute edge weights emanating from each transcription factor, representing direct transcriptional influence; and (2) eigenvector centrality, calculated using the igraph package (version 2.1.4), representing network-wide influence through connections to other highly connected nodes.

For each transcription factor, we calculated percentile ranks for both metrics across all TFs in the network, then computed an integrated master regulator score as the equally weighted average (50:50) of the two rank-normalized metrics. This weighting scheme was chosen based on the low correlation between the two metrics (Spearman ρ=0.146), ensuring they provide complementary information. Transcription factors were ranked by their master regulator scores, with the top 100 TFs retained for downstream comparative analysis. Specifically, for each transcription factor i in network N, the master regulator score was calculated as:

MR_score(i)=0.5×Percentile_rank(Weight_i)+0.5×Percentile_rank(EigenvectorCentrality_i)

where Weight_i = Σ|w_ij| for all target genes j regulated by TF i, and EigenvectorCentrality_i was computed using the eigen_centrality function from igraph on the directed regulatory network. Percentile ranks were calculated within each network independently, ranging from 0 (lowest) to 1 (highest). The resulting scores thus range from 0 to 1, with higher values indicating stronger combined regulatory influence.

### Cross-System Conservation Analysis

To identify master regulators conserved across reprogramming systems, we compared the top 100 ranked transcription factors from each system (MSC, adipocyte, human fibroblast). Within this top 100, we identified factors ranking in the top 30% (approximately top 30 TFs) across all three systems. Statistical significance of the observed overlap was assessed using permutation testing: we randomly shuffled transcription factor identities 10,000 times while preserving score distributions, calculated the expected overlap under the null hypothesis, and computed Z-scores and empirical p-values from the null distribution.

### Network Reorganization Analysis

To quantify changes in regulatory network architecture between baseline and reprogramming states, we calculated the following metrics for each transcription factor:

Effective out-degree: Calculated as the exponential of the Shannon entropy of the normalized regulatory weight distribution:

Deff=exp (−Σ pi log pi),

where pi=wi / Σ wj is the proportion of total regulatory weight allocated to target gene i. Higher values indicate more evenly distributed regulation across targets.

Gini coefficient: Measures inequality in regulatory weight distribution across targets, ranging from 0 (perfect equality) to 1 (maximum inequality). Lower values indicate more balanced regulation.

Regulatory coherence: For each transcription factor, we calculated the proportion of target genes where co-regulating TFs share the same regulatory direction (activation vs. repression). Coherence was computed as the mean across all targets, with higher values indicating greater regulatory alignment.

Pairwise TF correlations: Spearman correlations were calculated between regulatory weight vectors of all TF pairs using the cor function with method=“spearman”, yielding a correlation matrix for each network state.

Changes in these metrics (Δ) were computed as the difference between reprogramming and baseline states. Statistical significance was assessed using paired Wilcoxon signed-rank tests (wilcox.test with paired=TRUE, exact=FALSE to handle ties and zeros) for changes within each lineage, with n=9 core conserved transcription factors per system.

### Transcriptomic Age Analysis

Relative transcriptomic age for each cell was calculated using multi-tissue aging clocks trained to predict difference in chronological age and expected mortality (log hazard ratio).^46^ For mouse cells, we applied the mouse multi-tissue Bayesian Ridge (BR) clocks. For human fibroblasts, we applied the multi-species multi-tissue Bayesian Ridge clocks. Within each model, young control untreated cells were selected as a reference group, and all gene expression profiles were centered around median profiles of reference samples. tAge values were calculated across the pseudotime trajectory for all cells in each dataset. Correlations between master regulator expression and tAge were assessed using Spearman rank correlation. Correlations were calculated separately for young and aged cells within each system, with significance determined at p < 0.05.

### Ezh2 Perturbation Analysis

To establish causal relationships between master regulator activity and transcriptomic age, we analyzed publicly available gene expression data from Ezh2 perturbation experiments. For Ezh2 knockdown, we obtained data from siRNA-mediated inhibition in human dermal fibroblasts (GSE109064, n=3 biological replicates). For Ezh2 overexpression, we analyzed retinal ganglion cell data comparing wild-type Ezh2, catalytically inactive Ezh2-Y726D mutant, and control conditions (GSE247320, n=3 biological replicates per condition).

For each sample, tAge was calculated using both composite mortality and chronological aging clocks as described above. Mouse module-specific chronological and mortality multi-tissue clocks were applied to samples using the same methodology, followed by standardization of tAge across samples and calculation of average difference between the groups. P-values were assessed with one-tailed t-test and adjusted for multiple comparisons with the Benjamini-Hochberg approach. Differentially expressed genes were obtained from the original study, with genes considered significant at FDR < 0.05. Enrichment for rejuvenation signatures was assessed using hypergeometric tests. Directional concordance between Ezh2-perturbed genes and rejuvenation signatures was tested using Fisher’s exact test.

### Ezh2 Chromatin Accessibility Analysis

To understand transcriptomic differences between the Ezh2-WT and the catalytically inactive Ezh2-Y726D, we analyzed data of the differentially accessible regions (DAR) of each overexpression relative to their control conditions (n=3 biological replicates per condition) obtained from the Wang et al. 2024 paper.^61^ Comparison between the DARs and the rodent aging meta-signature were performed by subsetting any significant peaks (FDR<0.05, abs|LFC|>1) associated with genes that change with age (FDR <0.01, 7961 genes), and running Pearson correlation. To determine how genomic location impacts the association with age, we reran the analysis using peaks subset to specific classes (Distal Intergenic, Promoter, 5’ UTR, Exon, 3’UTR, Intron, or Downstream) and calculated the Pearson correlation by class.

Motif analysis was run separately for positive DARs (peak opening relative to the control) and negative DARs for Ezh2-WT and Ezh2-Y726D and performed using findMotifsGenome.pl from Homer (v5.1) with default parameters using a GC-matched background set. Comparison of motif enrichment results was performed using Spearman correlation between the ranked lists of motifs, ordered by p-value and filtered to motifs significant in at least one condition (FDR<0.001, 190 out of 490 computed motifs).

### Statistical Analysis and Reproducibility

Bootstrap confidence intervals (95%) for Spearman correlation coefficients were calculated using 10,000 bootstrap samples with replacement. For comparisons between young and aged correlations, we used Fisher’s z-transformation to test for significant differences between independent correlation coefficients.

All statistical tests were two-sided unless otherwise specified. P-values < 0.05 were considered statistically significant. Significance levels were denoted as: * p < 0.05, ** p < 0.01, *** p < 0.001, ns (not significant). Multiple testing correction was performed using the Benjamini-Hochberg FDR method where appropriate.

All analyses were performed in R version 4.5.1 using the following packages: NetID (version 0.1.1), Seurat (version 5.3.0), SingleCellExperiment (version 1.30.1), igraph (version 2.1.4), dplyr (version 1.1.4), ggplot2 (version 3.5.2), tidyr (version 1.3.1), patchwork (version 1.3.1), pheatmap (version 1.0.13), and cowplot (version 1.2.0).

## Data and materials availability

Additional data and materials are available online.

## Acknowledgements

Supported by NIA and Hevolution grants to VNG.

## Author Contributions

ADY, AT and VNG conceptualized the study. ADY performed all the analyses and wrote the first draft of the article. HPS contributed to the data analysis. VNG obtained funding. VNG and AT supervised the project. All authors contributed to the discussions and writing of the final draft.

