## Supplementary Figures for "Conserved Master Regulators Orchestrate Cellular Reprogramming-Induced Rejuvenation"

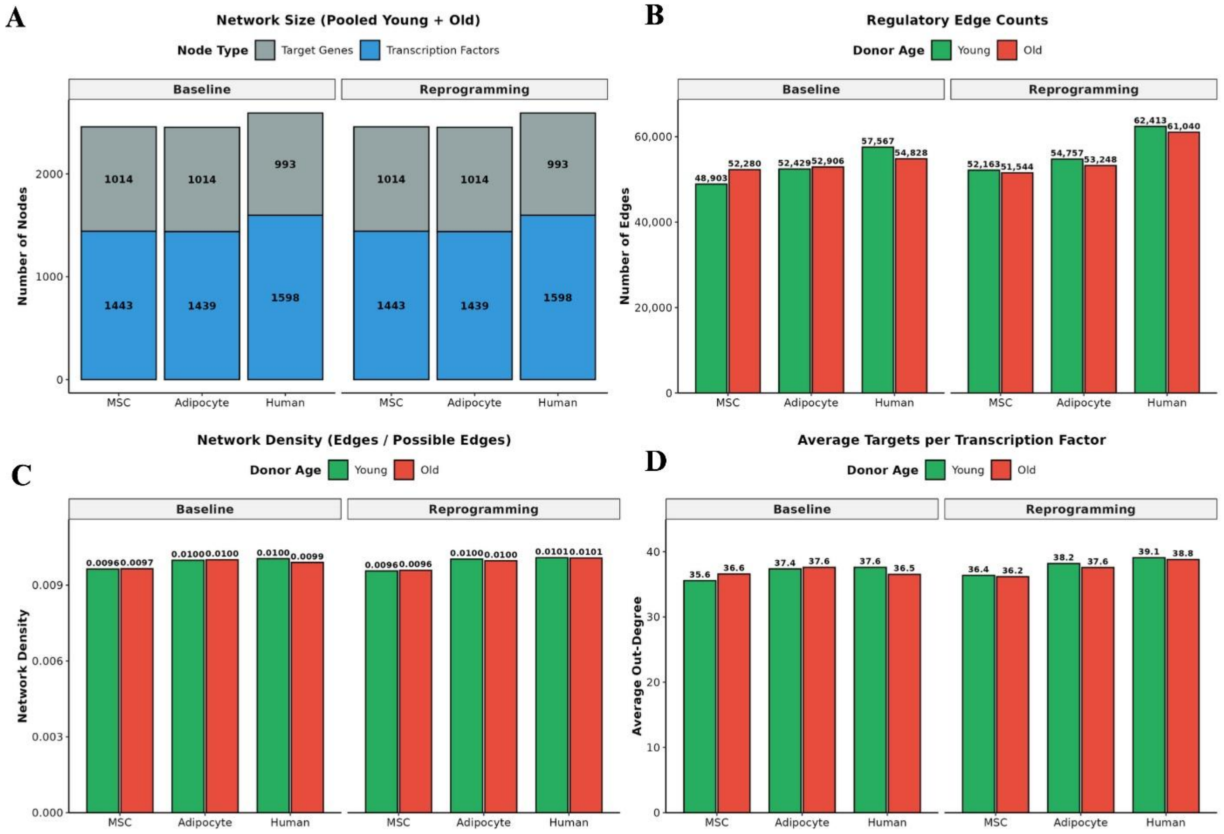


**Supplementary Figure 1. Basic network properties are consistent between young and old donors** (A) Network size showing the number of transcription factors (blue) and target genes (gray) in each reconstructed gene regulatory network. Networks were constructed using pooled young and old donor data. Node counts are similar across baseline and reprogramming states for all three cell types (MSC, Adipocyte, Human Fibroblast). (B) Regulatory edge counts stratified by donor age. Green bars represent young donors; red bars represent old donors. Edge counts are comparable between age groups within each cell type and condition, with similar numbers of regulatory interactions in young and old networks. (C) Network density calculated as the ratio of actual edges to possible edges. Networks show low density (~0.01) across all conditions, indicating sparse regulatory architectures typical of biological networks. Density values are highly similar between young (green) and old (red) donors. (D) Average out-degree (number of target genes per transcription factor). Values range from 35-40 targets per TF and remain consistent between young (green) and old (red) donors across all cell types and conditions.

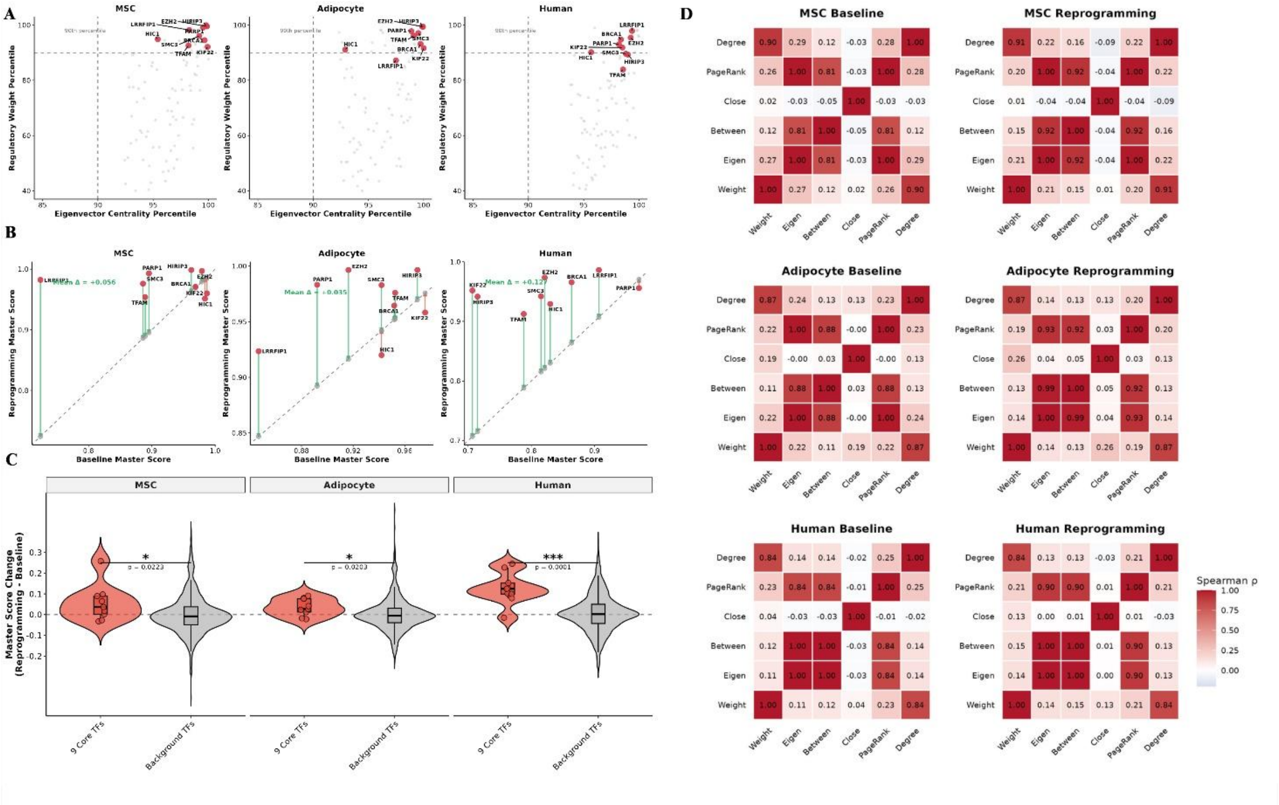


**Supplementary Figure 2. Master regulator identification integrates complementary network topology metrics** (A) The nine conserved master regulators score highly in both regulatory weight and eigenvector centrality. Scatter plots show regulatory weight percentile (y-axis) versus eigenvector centrality percentile (x-axis) for all transcription factors in each reprogramming system. Red points indicate the nine conserved factors (labeled); gray points indicate other TFs. Dashed lines mark the 80th percentile. All nine factors cluster in the upper right quadrant, demonstrating high scores in both dimensions across MSCs, adipocytes, and human fibroblasts. (B) Master regulator scores increase during reprogramming. Scatter plots compare baseline master scores (x-axis) versus reprogramming master scores (y-axis) for each system. Red points indicate the nine conserved factors; gray points indicate other TFs. The dashed line indicates identity (no change). Most factors show increased scores during reprogramming, with the nine conserved factors showing particularly robust increases. Mean score increase (Δ) is indicated for each system (MSC: $\Delta = +0.056$; Adipocyte: $\Delta = +0.035$; Human: $\Delta = +0.045$). (C) Master score increases are significantly enriched among conserved factors. Violin plots show the distribution of master score changes (Reprogramming - Baseline) for the nine conserved core TFs (red) versus all other background TFs (gray) in each system. The nine conserved factors show significantly larger score increases during reprogramming compared to background TFs (MSC: $p = 0.022$; Adipocyte: $p = 0.025$; Human: $p < 0.001$, Mann-Whitney U test). Box plots indicate median and quartiles; violin width indicates density. Statistical significance: * $p < 0.05$, *** $p < 0.001$. (D) Network centrality metrics provide complementary information. Correlation heatmaps show Spearman correlations between six network topology metrics (Degree, PageRank, Closeness, Betweenness, Eigenvector centrality, and regulatory Weight) for each system at baseline and during reprogramming. Color intensity represents correlation strength (red = positive, white = zero). Regulatory weight and eigenvector centrality show low correlation ($\rho \approx 0.1$-$0.3$), justifying their integration as complementary metrics. In contrast, PageRank, Betweenness, and Degree show high mutual correlations ($\rho > 0.8$), indicating redundancy. This supports the choice of weight and eigenvector centrality as the two non-redundant metrics for master regulator scoring.

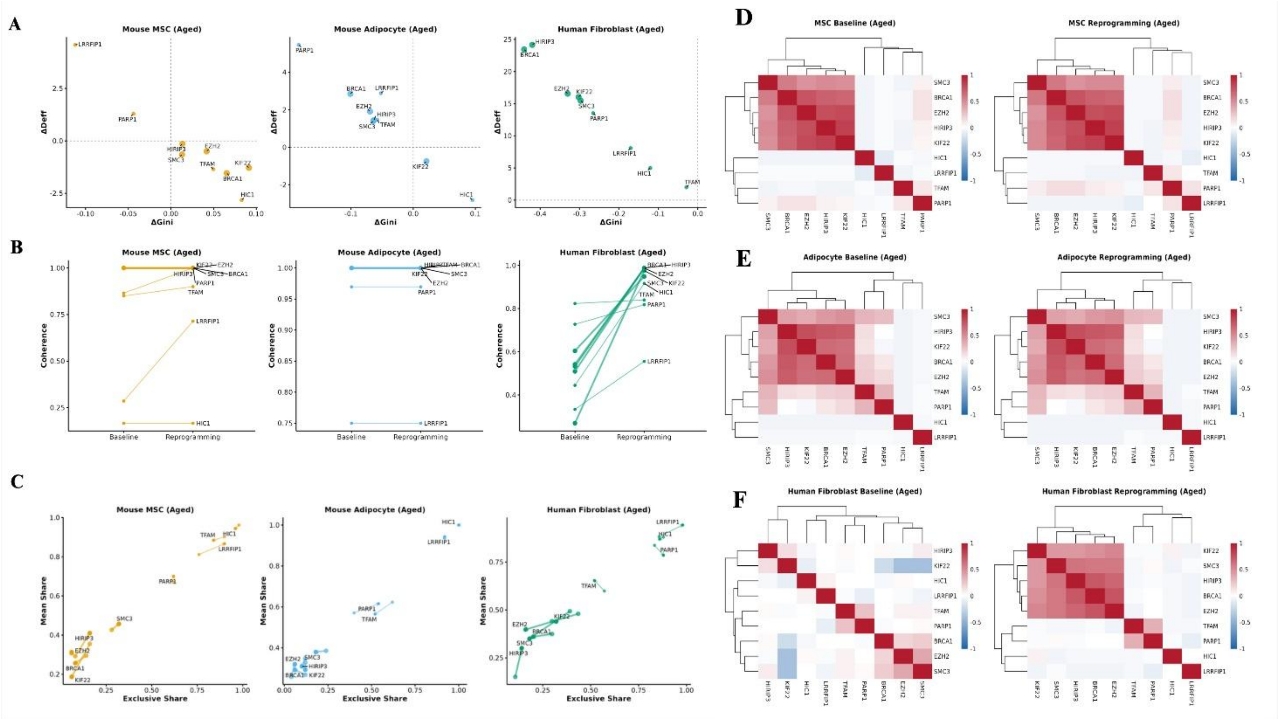


**Supplementary Figure 3. Network reorganization shows age-dependent differences in mouse but not human cells** (A) Changes during reprogramming in effective out-degree (ΔDeff) versus changes in Gini coefficient (ΔGini) for the nine conserved master regulators in old donors. Each point represents one transcription factor. MSC (Old): Factors show minimal or reversed changes compared to young donors, with several showing negative ΔDeff (network contraction). Adipocyte (Old): Factors show attenuated changes toward reweighting, with smaller magnitude shifts. Human Fibroblast (Old): Factors show robust increases in ΔDeff and decreases in ΔGini comparable to young donors, indicating preserved reorganization capacity. Dashed lines indicate no change. (B) Changes in regulatory coherence from baseline to reprogramming states in old donors. Lines connect the same transcription factor across states. MSC (Old): Minimal coherence increases, with baseline values already elevated (mean baseline = 0.796). Adipocyte (Old): Near-maximal baseline coherence (mean = 0.965) leaves little room for further increase. Human Fibroblast (Old): Clear coherence increases comparable to young donors (mean baseline = 0.532), demonstrating preserved network flexibility. (C) Regulatory weight distribution changes visualized as mean share versus exclusive share trajectories in old donors. Arrows show the transition from baseline (arrow origin) to reprogramming (arrowhead). MSC (Old): Disorganized trajectories indicating impaired reorganization. Adipocyte (Old): Minimal movement reflecting limited reorganization capacity. Human Fibroblast (Old): Consistent directional shifts toward distributed regulation, comparable to young donors. (D) Transcription factor regulatory weight correlation matrices for old donors at baseline (left) and during reprogramming (right). MSC (Old): Moderate baseline correlation (mean $\rho = 0.18$) with minimal change during reprogramming ($\Delta\rho = +0.01$), suggesting pre-existing network coordination. Adipocyte (Old): Similar baseline correlations (mean $\rho = 0.20$) with no net change during reprogramming ($\Delta\rho \approx 0$). Human Fibroblast (Old): Low baseline correlations (mean $\rho = 0.01$) with moderate increases during reprogramming ($\Delta\rho = +0.14$), demonstrating preserved network plasticity comparable to young donors. These patterns suggest that network reorganization capacity shows cell-type and species-specific variation with age, though the biological significance of these differences requires further investigation.
